# Macrophage migration inhibitory factor of Syrian golden hamster has similar structure and function as human MIF and promotes pancreatic tumor growth

**DOI:** 10.1101/449629

**Authors:** Pujarini Dash, Rajivgandhi Sundaram, Voddu Suresh, Surendra Chandra Sabat, Debasish Mohapatra, Sneha Mohanty, Dileep Vasudevan, Shantibhusan Senapati

**Affiliations:** Tumor Microenvironment and Animal Models Lab, Institute of Life Sciences, Bhubaneswar, Odisha, India; Macromolecular Crystallography Lab, Institute of Life Sciences, Bhubaneswar, Odisha, India; Manipal Academy of Higher Education, Manipal, Karnataka, India; Molecular Biology of Abiotic Stress Lab, Institute of Life Sciences, Bhubaneswar, Odisha, India; Department of Microbiology, Odisha University of Agriculture and Technology, Bhubaneswar, Odisha, India

**Author notes:** These authors have contributed equally to this work (Pujarini Dash, Rajivgandhi Sundaram, Voddu Suresh). **Address for correspondence:** Shantibhusan Senapati, BVSc & AH.; PhD, Scientist, Institute of Life Sciences, Nalco Square, Bhubaneswar-751023, Odisha, India., and Dileep Vasudevan, PhD, Scientist, Institute of Life Sciences, Nalco Square, Bhubaneswar-751023, Odisha, India.

## Abstract

Macrophage migration inhibitory factor (MIF) is a pleiotropic cytokine that increasingly is being studied in cancers and inflammatory diseases. Though murine models have been instrumental in understanding the functional role of MIF in different pathological conditions, the information obtained from these models is biased towards a specific species. In experimental science, results obtained from multiple clinically relevant animal models always provide convincing data that might recapitulate in humans. Syrian golden hamster (*Mesocricetus auratus*), is a clinically relevant animal model for multiple human diseases. Hence, the major objectives of this study were to characterize structure and function of hamster MIF, and finally evaluate its effect on pancreatic tumor growth *in vivo*. Initially, the recombinant hamster MIF (rha-MIF) was cloned, expressed and purified in bacterial expression system. The rha-MIF primary sequence, biochemical properties and crystal structure analysis showed a greater similarity with human MIF. The crystal structure of hamster MIF illustrates that it forms a homotrimer as known in human and mouse. However, hamster MIF exhibits some minor structural variations when compared to human and mouse MIF. The *in vitro* functional studies show that rha-MIF has tautomerase activity and enhances activation and migration of hamster peripheral blood mononuclear cells (PBMCs). Interestingly, injection of rha-MIF into HapT1 pancreatic tumor bearing hamsters significantly enhanced the tumor growth and tumor associated angiogenesis. Together, the current study shows a structural and functional similarity between hamster and human MIF. Moreover, it has demonstrated that a high-level of circulating MIF originating from non-tumor cells might also promote pancreatic tumor growth *in vivo*.

## Introduction

Clinically relevant animal models help in understanding the pathogenesis of different human and animal diseases and also play crucial roles in developing new therapeutics against them. In spite of the dominance of mouse as an experimental model animal, hamster has carved its niche as a potential model animal for studying many diseases and for evaluation of therapeutic agents. Syrian golden hamsters (*Mesocricetus auratus*) are frequently used in various disease pathogenesis studies due to the ease of handling them and the similarity to humans in disease development. Hamsters are important animal models for studying various infectious diseases of humans [1-8]. The hamster models of pancreatic and oral cancer have gained importance in their respective fields [9-14]. Moreover, these animals have also been instrumental in studying metabolic and/or inflammatory diseases like diabetes and pancreatitis [15, 16]. Despite the fact that Syrian golden hamster is an important clinically relevant animal model for different diseases, it is not used to its full potential. In this aspect, non-availability of complete genetic information of these animals and a lack of biological reagents related to them are the major constraints.

Macrophage migration inhibitory factor (MIF) is a pro-inflammatory cytokine with pleotropic functions in various pathophysiological processes [17-20]. Though initially identified as a T cell-derived cytokine that inhibits macrophage migration, its pleiotropic effects on immune cells, cancer cells, as well as non-cancerous cells made MIF more enigmatic to the researchers. Involvement of MIF in a number of human diseases like pulmonary hypertension, endothelial cell growth, atherosclerosis, wound healing, viral infection, many cancers including lung, colon, prostate, breast and pancreatic cancer has bagged a significant interest in this molecule [21]. In one of our earlier published study, we have characterized HapT1 cell line-based Syrian hamster tumor as a model of pancreatic cancer associated desmoplasia, an event which plays a key role in human pancreatic cancer progression [10]. In that study, for the first time, global proteomics analysis of whole cell lysate from hamster pancreatic stellate cells (PSCs) showed expression of macrophage migration inhibitory factor (MIF) by the cells [10]. At that particular time, due to unavailability of information and reagents for hamster MIF, we were unable to investigate the function of this molecule in the hamster model of pancreatic cancer. Hence, in the current study, our major objective was to characterize the hamster MIF protein, and evaluate the effect of exogenous MIF on the growth of pancreatic tumor in a syngeneic model of hamster pancreatic cancer.

In the current study, we have successfully purified recombinant hamster MIF protein (rha-MIF) from a bacterial protein expression system. Our analysis showed that like human MIF, rha-MIF also forms a trimer in solution. A commercially available MIF antibody raised against human MIF cross-reacts with rha-MIF. We resolved the trimeric rha-MIF crystal structure at 1.8 Å resolution, and the structural analysis showed multiple features in rha-MIF to be similar to mouse and human MIF. Further, biochemical and cell culture-based studies using endotoxin-free rha-MIF showed it’s enzymatic (tautomerase) and immunostimulatory activities, which suggest that the purified protein is biologically active. Importantly, all the biological properties of rha-MIF investigated in this study were similar to human MIF. At the end, we have investigated the effect of rha-MIF on the growth of HapT1 pancreatic tumor in its syngeneic host. The data clearly shows the pro-tumorigenic effect of rha-MIF on the HapT1 pancreatic tumor. Taken together, the data presented in this study have unraveled multiple information regarding hamster MIF, and indicate the importance of hamster as a model to investigate questions related to the role of MIF in pancreatic cancer progression.

## Materials and methods

### Recombinant hamster MIF (rha-MIF) expression, purification and Western blotting

Syrian golden hamster (*Mesocricetus auratus*) MIF open reading frame sequence (spanning residues 1-115) was PCR amplified and cloned in between NdeI and XhoI sites of a pET22b+ vector with an uncleavable C-terminal hexa-histidine tag. The protein was expressed in *E. coli* BL21 (DE3) cells at an OD_600nm_ of 0.6, by induction with 0.5 mM IPTG for 4 hours at 37 °C. Cells from 1 liter culture were pelleted down by centrifugation for 10 min at 7,000 r.p.m. and then suspended in 50 ml of buffer A containing 20 mM Tris-HCl (pH 7.5), 20 mM imidazole, 300 mM NaCl, 1 mM βME, 1 mM PMSF and one tablet of EDTA-free protease inhibitor cocktail (Sigma). The cells were lysed by sonication and the lysate was clarified by centrifugation at 18,000 r.p.m. for 45 min. Recombinant hamster MIF (rha-MIF) was first captured on a Ni-NTA affinity column (HisTrap FF 5 ml, GE Healthcare). Then the column was washed with 15 column volumes of buffer A and eluted with a linear gradient of buffer B (buffer A supplemented with 500 mM imidazole), followed by size-exclusion chromatography using a HiLoad 16/600 Superdex 75 pg column (GE Healthcare) with buffer C containing 20 mM Tris-HCl (pH 7.5), 150 mM NaCl and 1 mM DTT. The peak fractions containing MIF were pooled and concentrated to 30 mg/ml and stored at −80 °C. The purified protein was analyzed on 18% SDS-PAGE and stained with Coomassie Brilliant Blue to confirm the purity and to get an estimate of the monomeric molecular mass. To estimate the approximate molecular mass of purified hamster MIF in native conformation, *i.e*. to find whether the protein exists in an oligomeric form, analytical size-exclusion chromatography was performed using buffer C on a Superdex 200 10/300 GL column. Low molecular weight standards were used as a reference for molecular mass measurement. For biochemical experiments, endotoxin was removed from purified rha-MIF using Vivaspin endotest tubes (Sartorius Biotech, Germany). Next, the *E. coli*-derived recombinant hamster MIF used in our biochemical experiments was used to perform Western blot to elucidate the cross-reactivity of MIF polyclonal antibody (NBP1-81832; Novus Biologicals) towards hamster MIF, since this antibody was raised against a recombinant protein with 100% identity with human MIF and 89% identity with Syrian hamster MIF.

### CD spectroscopy

CD spectra of 10 μM rha-MIF in a buffer containing 10 mM sodium phosphate (pH 7.4) and 50 mM NaCl was recorded using a Chirascan CD spectroscope (Applied Photophysics). The spectra were collected in triplicate from 190 nm to 260 nm at 25°C at a bandwidth of 1.0 nm using a quartz cuvette with 10 mm path length. All the spectra were averaged and the buffer spectra was subtracted from those of sample. The estimation of secondary structural elements was done using the BeStSel server [22]. The CD intensities (in millidegree) are plotted against wavelength.

### Multiple sequence alignment and phylogenetic tree analysis

Protein sequences of Syrian golden hamster MIF (UniProtKB: A0A140EDM8), mouse MIF (UniProtKB: P34884) and human MIF (UniProtKB: P14174) were aligned using PRALINE multiple sequence alignment program and residue substitution matrix BLOSUM62 at http://www.ibi.vu.nl/programs/pralinewww/.PRALINE sequence conservation score 0 was given for the least conserved position and 10 for the most conserved position of alignment. Phylogenetic tree was constructed with MEGA 7 software [23] using the Neighbor-Joining method. [24]. The optimal tree with the sum of branch length is 0.15113287 and was drawn to scale with branch lengths (below the branches) in the same units as those of the evolutionary distances used to infer the phylogenetic tree. The evolutionary distances were computed using the Poisson method in the units of a number of residue substitutions per position.

### Crystallization and data collection

Recombinant hamster MIF (15 mg/ml) was screened for crystallization at 18 °C using commercially available screens with the help of an NT8 crystallization robot (Formulatrix, USA) in 96-well sitting drop crystallization plates. After 3 days, small and two-dimensional crystals appeared in a few conditions. The sequential micro-seeding approach was adopted for further optimization. Towards this end, crystals grown in a condition with 50% v/v PEG 400, 200 mM lithium sulphate and 100 mM sodium acetate (pH 7.5) were crushed and used for micro-seeding into the same condition, but in bigger drops with 1 μl protein and 1 μl condition, using hanging drop vapour diffusion method. After a week, bigger crystals appeared but were still two-dimensional in morphology. These crystals were picked, crushed and used as seed material for fresh crystallization screens following the random micro-seed matrix-screening method [25] in 96-well sitting drop plates, with the help of an NT8 crystallization robot. Crystals appeared in several new conditions; however, most of them were still two-dimensional and plate-like in morphology. A condition having 200 mM succinic acid (pH 7.0) and 20% w/v PEG 3350 yielded a bigger and thicker crystal after three weeks. The crystal was flash cooled in liquid nitrogen, after transferring to a solution with the composition of the crystallization condition, supplemented with 20% ethylene glycol. The diffraction images were recorded on an Eiger X 4M detector (DECTRIS) on beamline ID30A-3 of European Synchrotron Radiation Facility (Grenoble, France). 2000 images were collected with an oscillation of 0.1°, at a wavelength of 0.9677 Å.

### Data processing, structure determination and refinement

The diffraction images were processed using XDS [26]. The crystal structure of hamster MIF was solved by molecular replacement method using the program Molrep [27] from the CCP4 suite [28], using the mouse MIF crystal structure (PDB id: 1mff) as the search model. Crystallographic refinement and model building were performed iteratively using Refmac5 [29] and Coot [30]. The quality of the final model was assessed using the Ramachandran Plot from the program PROCHECK [31]. All the figures were prepared using PyMOL (Schrödinger, LLC). The data collection and refinement statistics are provided in **table 1**. The structure factors and refined coordinates have been deposited in the PDB, with the accession code 6ice.

**Table 1.** Crystal data collection and refinement statistics. Numbers in parentheses correspond to the last resolution shell.

| Parameter | Hamster MIF |
| --- | --- |
| **Data Collection** |  |
| Beamline | ESRF-ID30A-3 |
| Detector type | Eiger X 4M detector |
| Wavelength (Å) | 0.9677 |
| Data collection temperature (K) | 100 |
| Space group | P2<sub>1</sub>2<sub>1</sub>2<sub>1</sub> |
| a, b, c (Å) | 53.88, 55.85, 112.16 |
| $\alpha, \beta, \gamma$ (°) | 90, 90, 90 |
| Resolution (Å) | 38.85–1.84 (1.84–1.80) |
| R<sub>meas</sub> (%) | 0.118 (0.599) |
| I/$\sigma$(I) | 11.8 (3.5) |
| CC (1/2) (%) | 0.997 (0.868) |
| Total number of reflections | 239343(14289) |
| Mosaicity (°) | 0.65 |
| Completeness (%) | 98.5 (99.7) |
| R<sub>merge</sub> | 0.1 (0.559) |
| Multiplicity | 7.6 (7.8) |
| Wilson B-factor (Å<sup>2</sup>) | 15.8 |
| Matthews coefficient (Å<sup>3</sup>/Da) | 2.14 |
| Solvent content (%) | 42.4 |
| No. of molecules in ASU | 3 |
| **Refinement** |  |
| No. of unique reflections | 31590 (1840) |
| R<sub>work</sub>/R<sub>free</sub> (%) | 18.9/22.1 |
| Total no. of non-H atoms | 2790 |
| No. of water molecules | 211 |
| Mean B-factor (Å<sup>2</sup>) | 15.731 |
| **R.m.s. deviations** |  |
| Bond lengths (Å) | 0.0190 |
| Bond angles (°) | 1.8423 |
| **Ramachandran plot (%)** |  |
| Preferred region | 99.10 |
| Allowed region | 0.90 |
| Outliers | 0.0 |

### Tautomerase assay

MIF from other species has been reported to possess one unusual activity of catalyzing the tautomerization of D-dopachrome and L-dopachrome methyl ester into their corresponding indole derivatives. Tautomerase assay was carried out as described previously [32]. To analyze the tautomerase activity of rha-MIF, 4 mM of L-3,4-dihydroxyphenylalanine methyl ester was diluted in 5 ml autoclaved water and an appropriate amount of sodium periodate was added to it to make a final concentration of 8 mM. The mixture was incubated on ice for 20 min in the dark. 300 μl from this solution was mixed with 700 μl of the assay buffer consisting of 50 mM potassium phosphate and 1 mM EDTA (pH 6.0). Different concentrations of MIF were added to it and the decrease in absorbance at 475 nM was recorded after different time points of incubation.

### Effect of MIF on the expression of inflammation-associated genes in hamster peripheral blood mononuclear cells (ha-PBMCs)

To check the effect of MIF on the expression status of certain inflammation-associated genes in ha-PBMCs, cells were isolated from hamster blood using the Histopaque gradient as suggested by the manufacturer (Sigma, USA). The isolated ha-PBMCs were cultured in 10 % RPMI media and treated with rha-MIF (100 ng/ml), ISO-1 (100 μg/ml) or rha-MIF+ISO-1 for 4 hours. Then cells were harvested in RLT buffer and processed for RNA isolation as described by the manufacturer (Qiagen, USA). RNA quantification was carried out using Nanodrop followed by cDNA synthesis [10]. The expression levels of *Il-1β, Tnf-α, Vegf*, and *Il-6* were analyzed in all the cDNA samples through qPCR, by using gene-specific primers.

### Effect of MIF on PBMC migration

Prior to conducting any animal studies, all protocols were approved by the Institutional Animal Ethical Committee (Institute of Life Sciences, Bhubaneswar, India). Preparation of hamster peripheral blood mononuclear cells was carried out as previously described [33]. Briefly, normal hamsters having no symptomatic diseases were euthanized and blood was collected through the cardiac puncture in heparinized tube followed by Histopaque (Sigma, 11191) gradient centrifugation. After gradient centrifugation, the cells present in buffy coat layer were aspirated and washed twice with 10 ml PBS followed by resuspension in 2 to 3 ml of 0% FBS containing RPMI media. Cell counting was done through the trypan blue dye exclusion method. To check the effect of recombinant hamster MIF on the migration of hamster PBMCs, 3.5×10^5^ PBMCs in 0% FBS containing media (500 µl) were seeded in 8 µm pore trans-well insert (Millicell Hanging Cell Culture Insert, PET 8 µm, 12-well, MCEP12H48) and lower chamber was added either with 1% FBS containing media, 10% FBS containing media or 100 ng recombinant hamster MIF in 1% FBS containing media as chemoattractant (600 µl). Each experimental condition was carried out in triplicate. After 2 hours of incubation at 37 °C and 5% CO_2_, the cells were fixed in 10% buffered formalin for 10 min followed by staining with 0.1% crystal violet. Cells adhered to the upper surface of the membrane were removed with a cotton swab. Migrated cells at the lower surface of the insert were visualized under an inverted bright field microscope and 5 to 8 random images were captured for each membrane. The numbers of migrated cells were then expressed as the average of total field per membrane in triplicate for each experimental condition.

### Effect of rha-MIF on pancreatic tumor growth in vivo

Syrian golden hamsters (3-4 months old; male) were inoculated subcutaneously with 4×10^5^ HapT1 pancreatic cancer cells. Animals were checked for palpable tumors on daily basis. After six days from the injection of cancer cells, the tumor-bearing animals were randomized into two groups (n = 5 per group) and treated with rha-MIF (1 mg/kg) or PBS intraperitoneally for up to 13 days post inoculation of tumor cells. Tumor dimensions were measured every day, and tumor volume was calculated by using the formula V= (W^2^ X L)/2 for caliper measurements where V is the tumor volume, W is the tumor width and L is the tumor length. All the animals were sacrificed on 20^th^ day and tumors were harvested. After measuring the tumor weights, the samples were preserved and processed for histopathological analysis.

### Histology and immunohistochemistry

HapT1 tumors were dissected and fixed in 10% neutral buffered formalin at room temperature for 48 hours, and then tumor tissues were dehydrated and embedded in paraffin wax. 5 µm thick sections were cut and stained with hematoxylin-eosin. For immunohistochemistry, 5μm paraffin-embedded sections were de-paraffinized in xylene and rehydrated with graded ethanol and deionized water, then boiled in acidic pH citrate buffer (Vector Laboratories) for 20 min in a steam cooker. Endogenous peroxidase was blocked with 3% hydrogen peroxide in methanol for 20 min, and then washed with PBS followed by blocking with horse serum (Vector Lab) for 30 min. Then sections were incubated with the anti-Ki67 antibody (VP-RM04, Vector Laboratories) overnight at 4 °C in humidified chamber followed by washing with PBS. Then slides were incubated for 45 min at room temperature with a horse anti-rabbit/mouse IgG biotinylated universal antibody (Vector Laboratories) and then with ABC reagent for 30 min. The stain was developed with 3, 3′-diaminobenzidine (DAB; Vector Laboratories) substrate according to the manufacturer’s instructions. The slides were counter-stained with hematoxylin, dehydrated, cleared and mounted with mounting media (Vector Laboratories). Slides were observed under a Zeiss ApoTome.2 microscope and images captured at 10x, 20x and 40x magnifications. For quantification of blood vessels, multiple images from different areas of H&E stained HapT1 tumor sections were digitally captured and number of blood vessels/capillaries per field were counted.

### Effect of rha-MIF on proliferation of HapT1 cells

To check the effect of rha-MIF on the proliferation of HapT1 cells, 20,000 cells were seeded into the 24-well plate and allowed for attachment. Then, treatment with rha-MIF in different concentrations (0, 10, 50, 100, 150, 200, 300, 400 ng/ml) was given in triplicate to each group. After 48 hours, cells were washed twice with PBS and then fixed in 10% neutral buffered formalin for 10 min and stained with 0.6% crystal violet solution for 30 min, followed by de-staining with water. Quantification was done by dissolving crystal violet in 10% acetic acid for 15 min. Absorbance was measured at 470 nm using a Varioskan Flash Multimode Reader (Thermo Scientific).

### Effect of rha-MIF on VEGF expression in HapT1 cells

To check the effect of rha-MIF on the expression status of VEGF in HapT1 cells, 1,00,000 cells were seeded into 60 mm petri plate. After cells got attached, treatment was given to different groups such as control, rha-MIF (100 ng/ml) and rha-MIF+ISO-1 (100 µg/ml). After 48 hours of the treatment, cells were processed and RNA isolated using RNA isolation kit (Qiagen, USA). RNA quantification was done using Nanodrop spectrophotometer (Thermo Scientific) followed by cDNA synthesis. The expression status of *Vegf* was analyzed with the help of qPCR, by using the following specific primers: Forward: CTGGCTGGGTCACTAACA; Reverse: TTCTGGCTTTGTTCTGACTT.

## Results

### Expression and purification of Syrian golden hamster recombinant MIF (rha-MIF)

Although MIF was the first described cytokine, its role in some important pathophysiological conditions across different species has still remained a grey area. MIF from human and mouse or even from different human parasites have been structurally and functionally characterized [34-36]. In this study, we cloned and purified Syrian golden hamster (*Mesocricetus auratus*) MIF as described in materials and methods section. In one of our previous studies, we had cloned hamster MIF coding sequence (CDS) from m-RNA isolated from hamster pancreatic stellate cells [10]. Mouse genome is known to harbor multiple MIF-related sequences; however, only the true *Mif* gene has an intron/exon structure and a 5’ untranslated region [37]. Although the mouse MIF-related pseudogenes are highly homologous to cDNA, those contain different mutations that would generate truncated or altered MIF-like proteins. Hence, before cloning the hamster *Mif* CDS into the expression vector, we checked the hamster Mif genome organization. *In silico* analysis of recently updated genome sequence available in NCBI database (NW_004801628.1) confirmed that the Mif CDS cloned by us is from the true Mif gene, which has a similar genomic organization as human and mouse gene (**Supplementary Figure 1**). The expressed and purified rha-MIF showed an approximate molecular mass of about 12.5 kDa with more than 95% purity as observed from 18% SDS-PAGE, stained with Coomassie Brilliant Blue (**Figure 1A**). Comparison of the elution volume of rha-MIF with low molecular weight protein standards in the analytical size-exclusion chromatographic experiment suggested the molecular mass of rha-MIF in solution to be approximately three-times higher (39 kDa) than the monomer mass (12.5 kDa). This result indicates that the purified recombinant hamster MIF exists as a trimer (**Figure 1B**), like human and mouse MIF. CD spectroscopic analysis showed the recombinant protein to be properly folded. The secondary structure content of the protein was estimated to be 27 % α-helix, 54 % β-sheet, 8.8 % turns and 10.2 % unordered structure, with an RMSD and NMRSD values of 0.40 and 0.0237, respectively, as given in the form of a pie chart along with the profile (**Figure 1C**). The Western blot analysis revealed the cross-reactivity of MIF polyclonal antibody (NBP1-81832; Novus Biologicals) towards hamster MIF and further confirmed successful expression and purification of the protein (**Figure 1D**).

**Figure 1.**
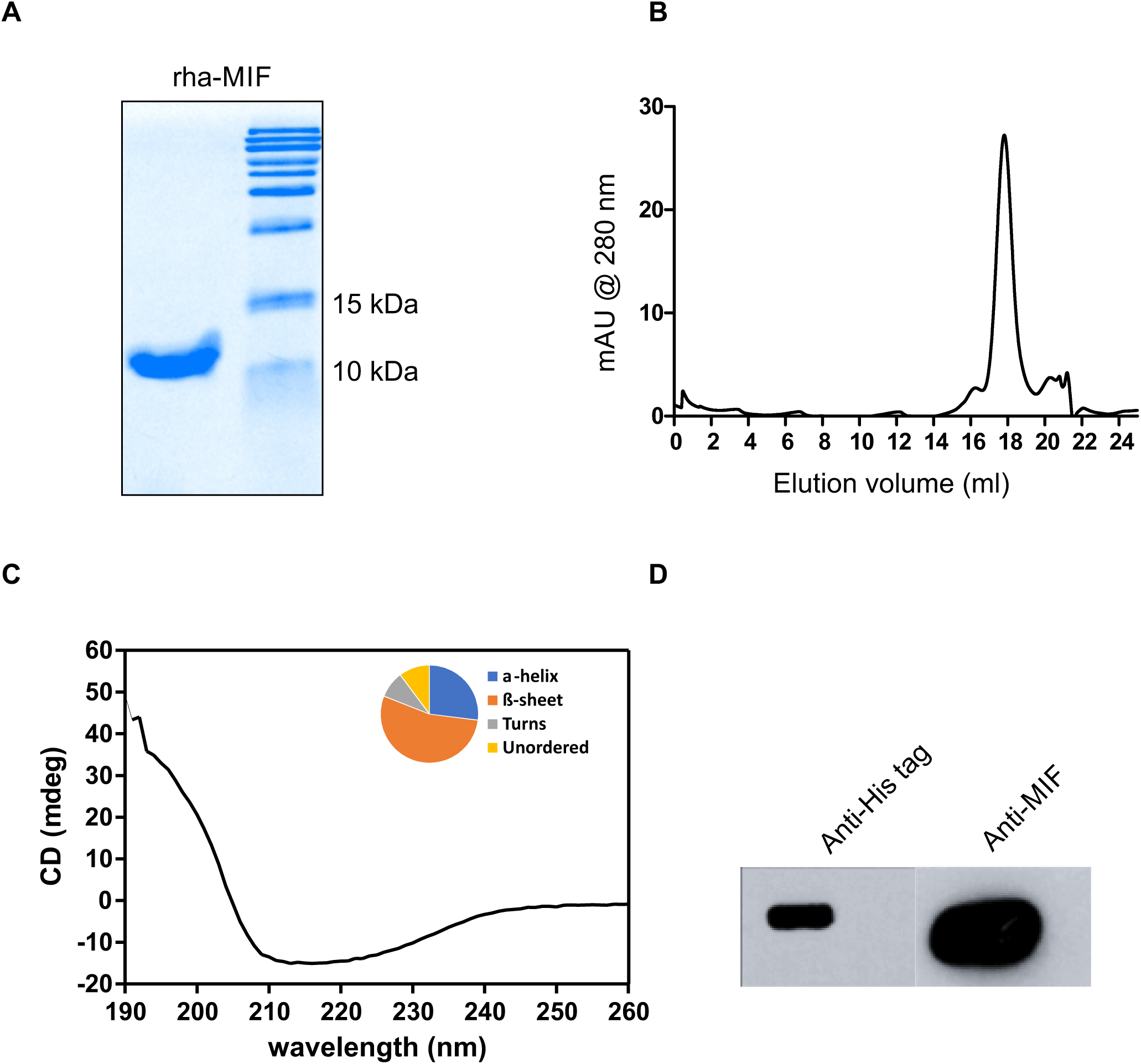
Expression and purification of rha-MIF. (A) SDS-PAGE analysis of purified rha-MIF suggesting an approximate molecular mass of ∼12.5 kDa for the monomer. (B) Superdex 200 10/300 GL analytical size-exclusion chromatography profile of rha-MIF suggesting an apparent molecular mass of ∼39 kDa in solution, corresponding to the size of a trimeric form. (C) CD spectrum of rha-MIF confirming a properly folded protein. The pie chart in the inset shows the secondary structure make-up of the protein. (D) Western blot analyses with polyclonal anti-His and anti-MIF antibodies confirming the expressed protein to be rha-MIF.

### Sequence and phylogenetic analysis of ha-MIF

Alignment of the primary sequences of MIF from hamster, mouse, and human were performed with PRALINE program. **Figure 2A** indicates that hamster MIF shares 93 % and 89 % sequence identity with mouse MIF and human MIF, respectively. The alignment reveals that the residues responsible for tautomerase activity and substrate binding such as Pro2, Lys33, Ile65, Tyr96 and Asn98 are conserved in hamster MIF as well. In addition, the ‘CALC’ motif required for oxidoreductase activity is also conserved in hamster MIF (**Figure 2A**). As both the tautomerase and oxidoreductase sites are present in hamster MIF, it might have similar enzymatic and biological activities as that of human and mouse MIF. The phylogenetic analysis of MIF from hamster, mouse and human was carried out using the Neighbor-Joining method with the help of MEGA 7 software further confirms that hamster MIF is evolutionarily closer to the mouse MIF than human MIF (**Figure 2B**).

**Figure 2.**
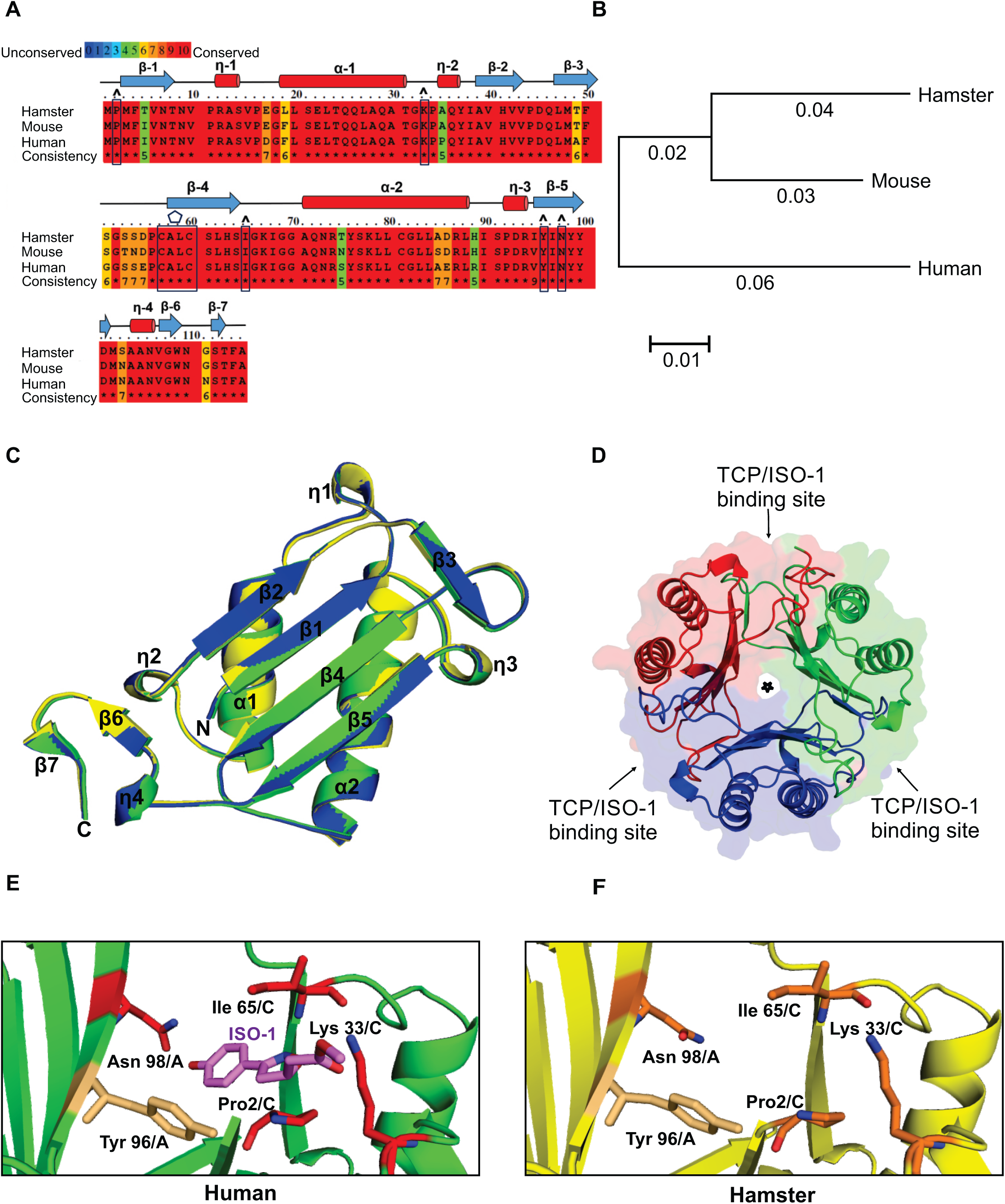
Syrian golden hamster MIF protein sequence and structural analysis. (A) Primary sequences of MIF from Syrian golden hamster (UniProt accession number A0A140EDM8) mouse (P34884) and human (P14174) were aligned with PRALINE program. The conserved residues are shown in red color. The boxes with cap indicate the conserved residues of the tautomerase active site, and the polygon label above the boxed sequence shows the conserved ‘CALC’ motif responsible for the oxidoreductase activity. Secondary structural elements of MIF are indicated above the aligned sequences. (B) Phylogenetic tree of hamster, mouse and human MIF constructed using MEGA 7.0 software. The branch-lengths indicate the evolutionary distances. (C) Superposition of the monomeric structure of MIF from hamster (yellow; PDB id: 6ice, from this work), mouse (blue; PDB id: 1mfi) and human (green; PDB id: 3djh) in cartoon representation, revealing similar 3D-structure. The hamster MIF structure aligns with human MIF and mouse MIF with RMSDs of 0.24 Å and 0.26 Å, respectively for the Cα atoms. (D) Crystal structure of homo-trimeric hamster MIF in cartoon representation, along with semi-transparent surface representation. The monomers are colored in red, green and blue. The three tautomerase catalytic pockets (TCP; also, the sites for ISO-1 binding) located at the monomer-monomer interfaces are labelled with arrows and the central solvent-accessible channel is marked with a star sign. (E) A zoomed view of the tautomerase catalytic pocket (TCP) of the ISO-1 bound human MIF structure (PDB id: 1ljt) in green, wherein the active site residues Pro2, Lys33, Ile65 and Asn98 that are interacting with ISO-1 are colored in red and Tyr96 which is part of the catalytic pocket, but not involved in ISO-1 interaction is colored in light orange; all in stick representation. ISO-1 is shown in stick representation, in magenta color. (F) A similar zoomed view as (E) for hamster MIF TCP from this work, wherein the structure is shown in yellow, the active site residues Pro2, Lys33, Ile65 and Asn98 are shown in orange and Tyr96 in light orange; all in stick representation.

### Crystal Structure of hamster rha-MIF

The rha-MIF crystal belonged to the orthorhombic P2_1_2_1_2_1_ space group with a = 53.88 Å, b = 55.85 Å, c = 112.16 Å, α=β=γ=90° as unit cell parameters. X-ray diffraction data collected at ESRF-ID30A-3 beamline was used to solve the structure. The data collection and refinement statistics are given in **table 1**. The crystal structure of rha-MIF was resolved to 1.8 Å resolution. It forms a trimer, with three identical molecules in an asymmetric unit. A monomer of rha-MIF is made up of six beta strands, four of which form a mixed beta sheet within one subunit, two antiparallel alpha helices and two shorter beta strands on either end of the sheet (**Figure 2C**). Also, a very small beta strand (β7) is present towards the C-terminus. The monomer structure is highly conserved among MIF from human, mouse and hamster, as could be observed from the structure alignment (**Figure 2C**). The RMSD for the Cα backbone atoms between hamster and human MIF (PDB id: 3djh) and between hamster and mouse MIF (PDB id: 1mfi) are 0.24 Å and 0.26 Å, respectively. The trimer adopts a barrel-like architecture formed by three inter-connected monomeric subunits (**Figure 2D**), as has been reported for other MIF structures. The trimeric rha-MIF structure is retained by the short beta strands β3 of one monomer and β6 of the third monomer becoming part of the beta sheet formed by the beta strands β1, β2, β4 and β5 of the second monomer in between, together forming a six-stranded beta sheet. Herein, β3 of monomer-1 comes in close proximity and form hydrogen-bond network, typical of beta sheets, with β2 of the monomer-2 and β6 of monomer-3 comes close to β5 of monomer-2. This contribution of beta strands by all three monomers to their respective adjacent monomers to form beta sheets seems to provide stability to the trimeric structure. In the trimeric ring structure of MIF, the beta sheets of the three monomers together form a solvent-accessible central water channel, whose role in the context of MIF function is still not known [34]. A MIF trimer has been reported to have three catalytic pockets for tautomerase activity, located in the monomer-monomer interface between the three monomers (**Figure 2D**). The residues Pro2, Lys33 and Ile65 from one monomer and the residues Tyr96’ and Asn98’ from its adjacent monomer [38], together form the catalytic pocket responsible for the tautomerase activity in human and mouse MIF and these residues are conserved in hamster MIF as well. This would explain why rha-MIF also shows tautomerase activity (described in the next section). Tautomerase activity of human MIF is known to be inhibited by the MIF-inhibitor ISO-1 [39]. ISO-1 binds to the residues Pro2, Lys33, Ile65 and Asn98’ [40], all of which form part of the tautomerase catalytic site located at the interface of two monomers (**Figure 2E; Supplementary Figure 2**). The Tyr96 residue which is a part of the catalytic pocket is however not involved in ISO-1 binding. As the residues of the catalytic pocket and ISO-1 binding are completely conserved in hamster MIF (**Figure 2F**), ISO-1 and other similar compounds binding to the catalytic pocket would block hamster MIF catalytic function as well. Though previously reported human and mouse MIF structures had sulfate ions forming part of the crystal structure [41-43], hamster MIF structure from this work did not have any sulfate ions, presumably because the final crystallization condition did not have any sulfate.

### Tautomerase activity of rha-MIF

Unlike all other cytokines, MIF has a tautomerase activity, and it performs this enzymatic function through an N-terminal catalytic proline base [44]. In this study, the tautomerase activity of the purified rha-MIF was analyzed using L-dopachrome methyl ester substrate. The activity was determined after background correction for the non-enzymatic activity (dopachrome dependent non-specific activity; **Supplementary Figure 3**). As the linear range of the kinetic assay for the L-dopachrome methyl ester is too short, the slope value from the initial linear part (ranging from 0 to 15 sec.) of the declining curve (Δ abs. s ^-1^) was considered to determine the activity (**Supplementary Figure 3**). The rate of tautomerase activity of the protein (*i.e*., the amount of the product formed per second) showed a typical enzymatic response, wherein activity increased with increasing concentration of protein at lower concentrations and revealed a saturation tendency at higher concentrations (**Figure 3A**). The initial velocities of the reaction were measured as a function of substrate concentration from 0 to 800 μM (nmol ml^-1^). The data presented in **Figure 3B** was used to determine the K_m_ by non-linear least squares fit method following simple Michaelis-Menten equation (**Figure 3C**). The calculated K_m_ value is 665 μM (nmol ml^-1^).

**Figure 3.**
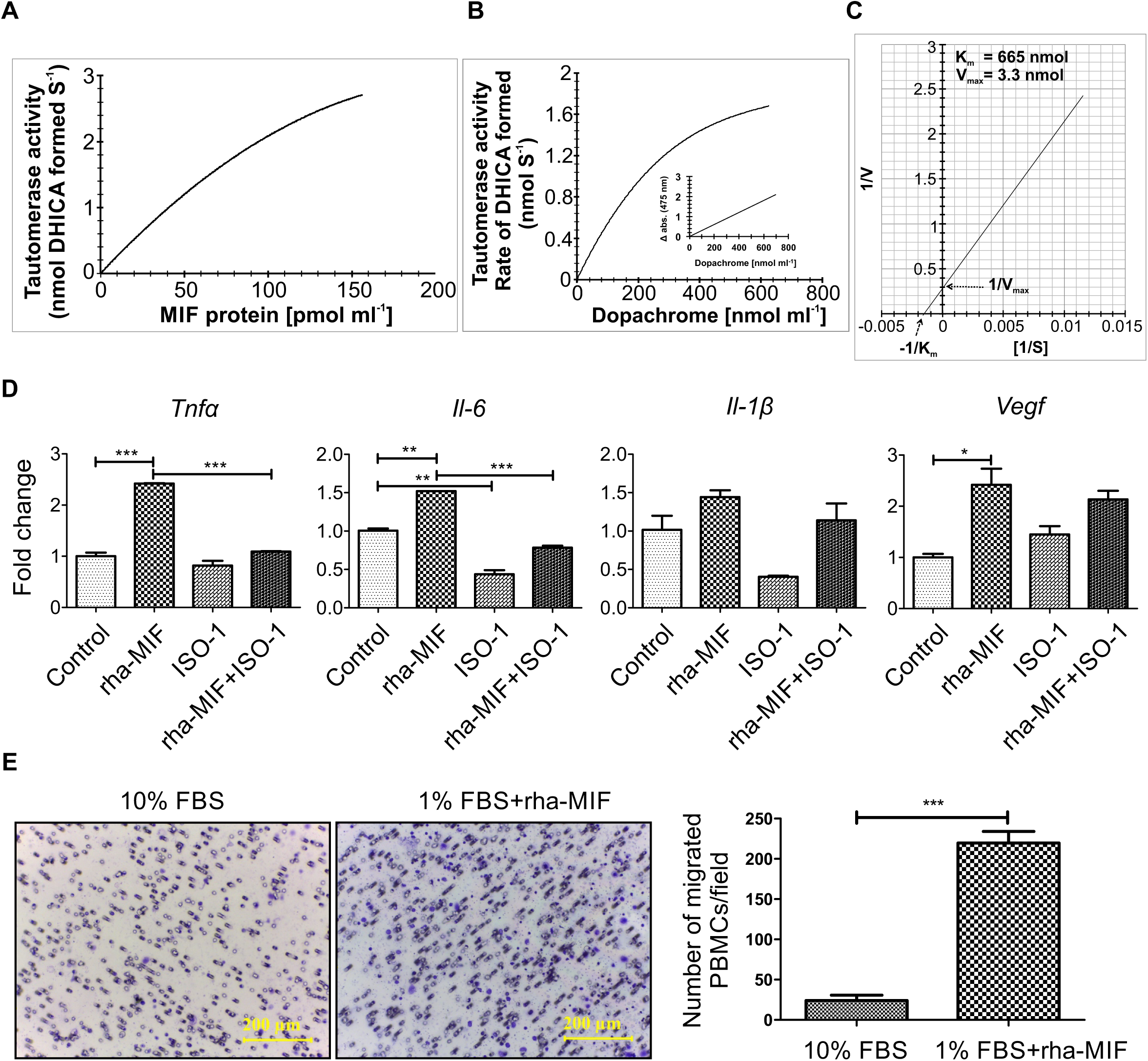
*In vitro* functional analysis of rha-MIF. (A) Data shows the rha-MIF-tautomerase activity under increasing concentration of this protein. (B) Graphical representation showing influence of increasing concentrations of dopachrome on MIF-tautomerization activity (C) Determination the K_m_ and V_max_ for the substrate–dependent MIF-tautomerase activity. (D) Figure shows the expression of *Tnf-*α, *Il-6, Il-1*β and *Vegf* in ha-PBMCs following stimulation with rha-MIF (100 ng/ml), ISO-1 (100 μg/ml) and rha-MIF+ISO-1 for 4 hours. Data presented as mean ± S.D., *p < 0.05; **p < 0.005; ***p < 0.0005 using one-way ANOVA with Bonferroni’s Multiple Comparison Test. (E) Results of the trans-well migration assay show the effect of rha-MIF on migration property of haPBMCs. Digital images of the stained trans-well porous membrane show PBMCs (blue colored) that have migrated through the pores (in presence and absence of rha-MIF). Quantification of number of migrated cells in different conditions is shown through bar graph. Each bar represents the mean ± SEM. ***p < 0.0005 using a Student t test.

### Effect of rha-MIF on activation and migration of PBMCs

MIF is known as a potent pro-inflammatory cytokine and regulates activation and migration of immune cells [45]. Exogenous MIF treatment induces expression of various pro-inflammatory cytokines (e.g. TNF-α, IL-6, IL-1β, and IL-8) and angiogenic factors (e.g. VEGF) in different immune and/or endothelial cells [46-48]. In this study, we investigated the effect of endotoxin-free rha-MIF on the activation of hamster PBMCs (haPBMCs) in the presence and absence of ISO-1. As readout of PBMC activation, the level of *Tnf-α, Il-6, Il-1β*, and *Vegf* expression was measured through quantitative PCR analysis. Exposure to exogenous rha-MIF induced significantly higher expression (p<0.0005) of *Tnf-α, Il-6*, and *Vegf* in haPBMCs (**Figure 3D**). Except for *Vegf*, the rha-MIF-induced expression of other molecules (*Il-1β, Il-6, Tnf-α*) got considerably suppressed upon ISO-1 treatment (**Figure 3D**). Interestingly, ISO-1 itself suppressed the endogenous level of *Il-1β* and *Il-6* in haPBMCs, which indicates the role of endogenous MIF in regulating expression of these molecules in PBMCs (**Figure 3D**). Effect of different cytokines on immune cell activation and migration/recruitment together contribute to the overall inflammation at the site of injury and/or infection. Human MIF is known to act as a chemoattractant for PBMCs [45]. Hence, we wanted to check whether rha-MIF has a similar effect on the migration properties of haPBMCs. The results of our trans-well migration assay showed significantly higher number of migrated PBMCs (p<0.0001) in conditions where rha-MIF was used as a chemoattractant than the control (**Figure 3E**).

### Effect of rha-MIF on the growth of HapT1 pancreatic tumor in its syngeneic host

In pancreatic cancer patients, the tumor tissues and circulating blood have a higher level of MIF than in healthy subjects [49, 50]. Pancreatic cancer cells overexpress MIF and are believed to be the major contributor to the overall MIF level in the pancreatic tumor microenvironment. At the same time, multiple normal cells like immune cells, mesenchymal stem cells, endothelial cells, etc. also express MIF [51]. To understand the pathophysiological role of MIF in pancreatic cancer progression, most of the studies have adapted genetic and chemical targeting of MIF expressed by the cancer cells. However, in these approaches, the effect of MIF produced by normal cells on cancer cells (paracrine effects) is not being addressed. Hence, in the current study, we evaluated the impact of rha-MIF on the growth of HapT1 tumors in its syngeneic host. The data in **Figures 4A and 4B** shows that systemic administration of rha-MIF significantly enhanced HapT1 tumor growth. Though tumor uptake was 100% in both PBS and rha-MIF injected groups, but after 20 days of cancer cells injection, the tumor weight of rha-MIF injected tumors was significantly higher (p<0.0005) than PBS injected tumors (**Figure 4C**). Moreover, the presence of a substantially higher number of Ki67-positive cells (p<0.0001) in the tumors of rha-MIF-injected animals than PBS injected animals (**Figures 4D and 4E**) corroborates the observed pro-tumorigenic effect of exogenous MIF on pancreatic tumors *in vivo*.

**Figure 4.**
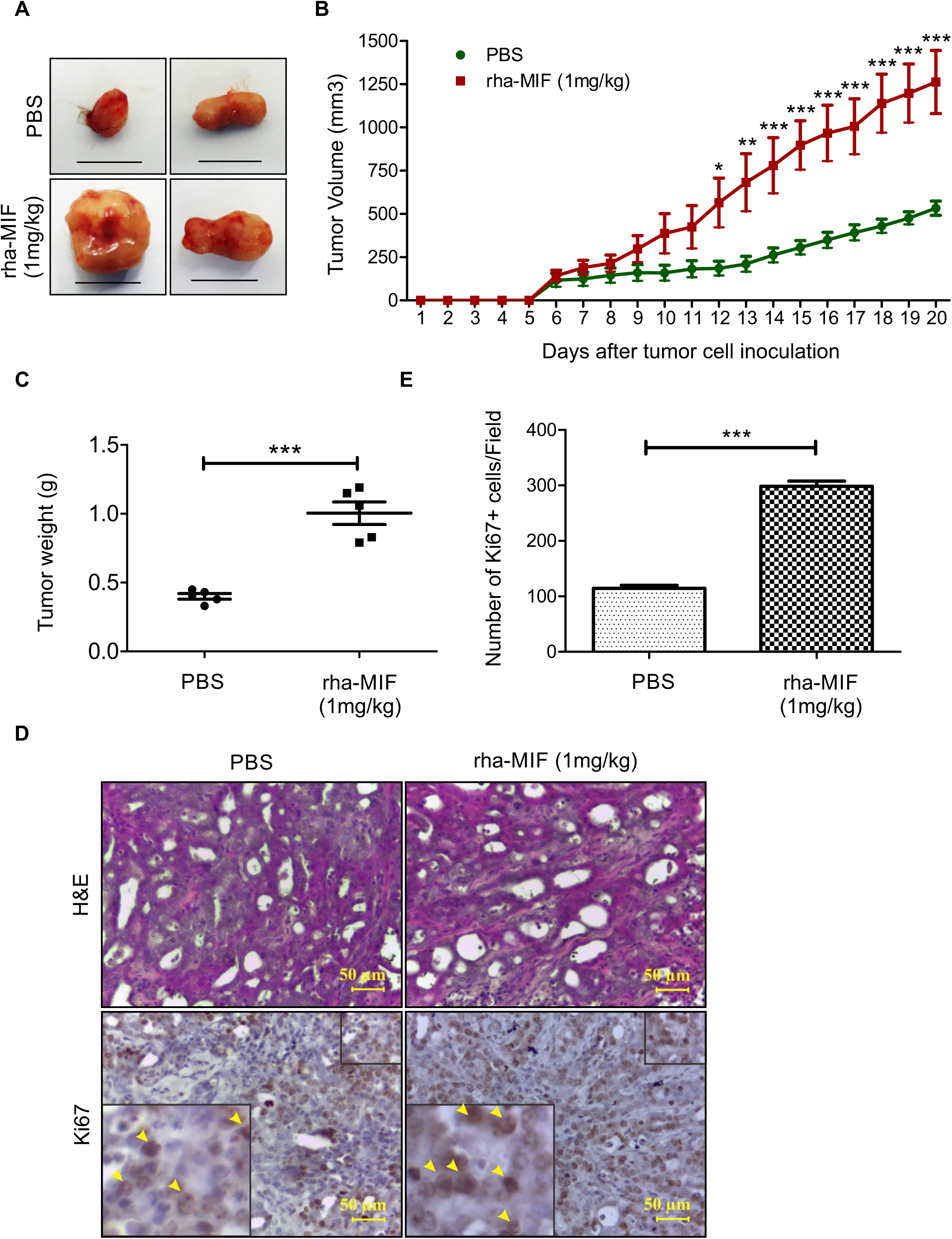
Effect of rha-MIF on HapT1 tumor growth in its syngeneic host. (A) Representative digital images of HapT1 tumors obtained from PBS or rha-MIF injected animals show bigger tumor mass in rha-MIF injected groups. The scale bar is equivalent to 1cm (B) Graphical representation of tumor volume calculated every day after initiation of treatment revealed significant increase in the tumor growth in rha-MIF injected animals. Data represents the mean ± SEM for five hamsters per group. ***p < 0.0001 using two-way ANOVA (C) Dot-plot graph showing significant increase in HapT1 tumor weight in rha-MIF treated animals than control. Data represents the mean ± SEM for five hamsters per group. ***p < 0.0001 using a Student t test. (D) Hematoxylin and Eosin (H&E) stained HapT1 tumor tissue sections shows moderately differentiated tumors in all the conditions (PBS or rha-MIF injected). The (A) lower panel representative images show tissue sections stained with Ki67 antibody. Images were captured at 40× magnification. (E) Graph showing number of Ki67 positive cells/field. Each bar represents the mean ± SEM for five hamsters per group. ***p < 0.0001 using a Student t test.

### Mechanistic aspects of rha-MIF-mediated HapT1 tumor growth in its syngeneic host

Overexpression of MIF by cancer cells is known to promote tumor growth through multiple mechanisms [52]. To check the direct effect of rha-MIF on HapT1 cancer cells growth, we treated HapT1 cancer cells with rha-MIF *in vitro* and after 2 days, the number of viable cells was quantified through crystal violate staining of the cultured cells. The data shown in **Figure 5A** shows that exogenous rha-MIF does not have a significant effect on the overall growth of HapT1 cells *in vitro*. This result promoted us to check the status of MIF receptor expression in HapT1 cells, and our quantitative PCR analysis data clearly shows high level of Cd74 and low level of Cxcr4, and Cxcr2 expression in HapT1 cells (**Figure 5B**). At the molecular level, to check the response of HapT1 cells to exogenously treated rha-MIF, we checked the status *Vegf* expression in rha-MIF-treated and -untreated HapT1 cells. The gene expression analysis showed a significant up regulation in *Vegf* expression upon rha-MIF treatment (**Figure 5C**). MIF is known to induce tumor angiogenesis in different cancers [53]. Moreover, up regulation of *Vegf* in HapT1 cells upon rha-MIF stimulation suggests that rha-MIF might have an effect of HapT1 tumor angiogenesis. To check this possibility, HapT1 tumor tissues treated with or without rha-MIF were microscopically analyzed and the number of blood vessels was quantified. Quantification of vessel density clearly showed significantly higher number of blood vessels in rha-MIF treated tumors than control tumors (**Figure 5C**). Together, the data suggest that circulating MIF of non-tumor origin might promote tumor angiogenesis, thereby promoting overall tumor growth *in vivo*.

**Figure 5.**
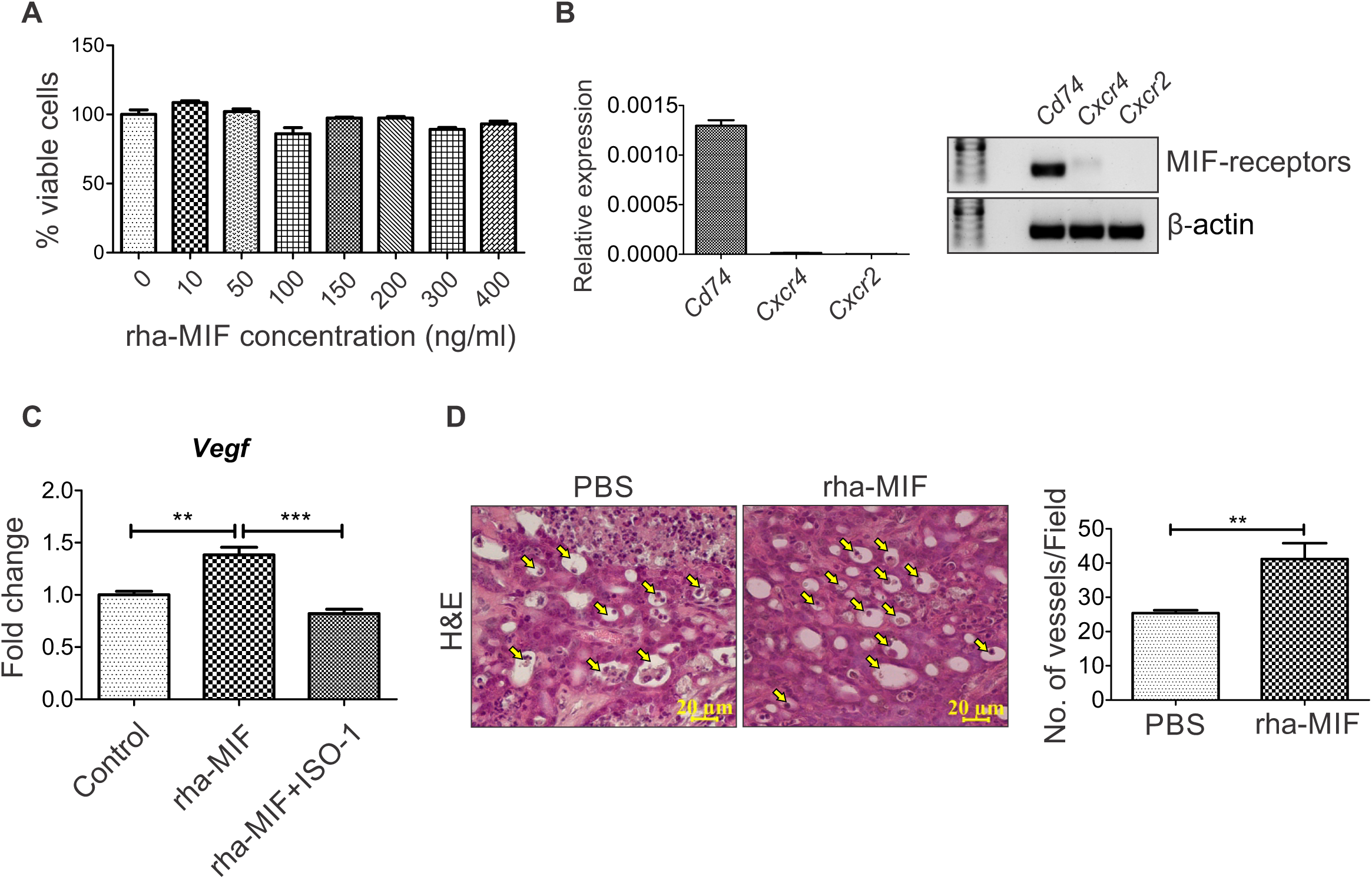
Effect of rha-MIF on HapT1 cells viability and angiogenesis. (A) Graph showing the percentage viable cells upon treatment with different concentrations of rha-MIF after 48 hours. (B) Graph showing the relative expression of *Cd74, Cxcr4* and *Cxcr2* in HapT1 cells and agarose gel image showing the amplified products in qPCR. Relative expression is calculated using formula 2^-dCT^, ΔCT= CT_gene_– CT_β-actin_. (C) Graph showing the expression of *Vegf* in HapT1 cells following stimulation with rha-MIF (100 ng/ml) and rha-MIF+ISO-1 (100 µg/ml) for 48 hours. Each bar represents the mean ± SEM. ***p < 0.005 using one-way ANOVA with Bonferroni’s Multiple Comparison Test. (D) Representative H&E images (captured in 40x magnification) showing the number of blood vessels (arrows indicated) between PBS and rha-MIF treated tumor tissues. Quantification was done by counting the number of vessels/field in 20x magnification images. Each bar represents the mean ± the SEM. **p < 0.05 using a Student t test.

## Discussion

The cross-reactivity of MIF polyclonal antibody (NBP1-81832; Novus Biologicals) generated against a recombinant human MIF having 89% identity with Syrian hamster MIF indicates conserved antigenic epitopes in human and hamster MIF. Moreover, these results confirm the reliability of using NBP1-81832 antibody to detect Syrian golden hamster MIF. Primary sequence and crystal structure analysis of rha-MIF showed that mouse and hamster have almost similar level of identities with human MIF, which indicates that experiments related to MIF in mouse and hamster might have also similar outcomes. However, it will be apt to mention that the overall outcome of an experimental intervention in any animal model depends on multiple factors like the animal’s genetic makeup, physiology, natural behavior, etc. Hence, in certain instances, hamster might be a more clinically relevant model than mice and *vice versa*. For example, hamster is a good host for multiple infectious agents to which mouse are resistant [54]. To investigate the role of MIF in the pathogenesis of these diseases, hamster model might be instrumental. Moreover, conserved structural features like the active site for tautomerase and ISO-1 binding in hamster MIF further justifies the suitability and relevance of this model in MIF-related studies (**Figures 2D and 2E; Supplementary Figure 2**). A study has shown that the diameter of the central channel of MIF from parasites like *Giardia lamblia* and *Plasmodium* species is narrower than that of human MIF. This structural difference between human- and parasite-MIF central channels might form a basis for structure-based drug design against the parasites [55]. A wider channel size in human- and hamster-MIF (**Figure 2D**) indicates that this structure might have a functional significance in vertebrates, which warrants further investigation.

The K_m_ value of rha-MIF protein (**Figure 3C**) is not in agreement with the MIF proteins of other species (2.4 mM: [56]; 2.7 mM: [57]; 2.1 mM: [58]). This enzyme, was found to have much superior activity, with nearly 4-fold higher sensitivity towards L-dopachrome. This difference might be due to an itrinsic superior enzymatic activity of hamster MIF and/or difference associated with the technical process (protein quality, instrument sensitity, etc.). A further in-depth investigation to compare the tautomerase activity of hamster MIF and human MIF, using the same techical procedure might be helpful while exploring for effective MIF-inhibitors. Being a conserved protein and having a unique combination of hormone-, cytokine- and thioredoxin-like properties, MIF is considered as a potent cytokine with pleiotropic effects on immune and inflammatory events [59]. It is known to regulate expression of pro-inflammatory mediators leading to early patient death in sepsis [60, 61]. At the same time, human MIF is known to have differential effect on the migratory properties of various cell types [45]. The recruitment of immune cells to the site of infection leads to clearance of viral, bacterial and fungal infection but on the other hand, monocytes also contribute to their pathogenesis and inflammation [62]. Effect of hamster MIF in augmenting the directional migration of hamster PBMCs towards it shows its chemo-attractant property similar to human MIF (**Figure 3E**) [45]. Moreover, the effect of ISO-1 in inhibiting the rha-MIF mediated upregulation of the expression of proinflammatory molecules and migration of hamster PBMCs corroborates the information obtained from the structural analysis of rha-MIF. These data further suggest that, ISO-1 could be reliably used as a ha-MIF inhibitor in various *in vitro* and *in vivo* studies.

Due to its remarkable involvement in cytokine cascade in tumor microenvironment, MIF is described as the connecting link bridging cancer with inflammation [63] and it is also known to be implicated in angiogenesis in multiple cancer types [64-66]. In recent past, studies have demonstrated the presence of high level of MIF in different cancer patients. A high level of MIF is reported from the tumor tissues and circulating blood of pancreatic ductal adenocarcinoma (PDAC) patients [49, 50]. Expression level of MIF correlates with poor prognosis of PDAC patients. In most of the studies trying to understand the role of MIF in the progression of different cancers, MIF expression was knocked down in overexpressing cancer cells. However, these attempts do not address the contribution of MIF originating from stromal cells of cancer patients. To best of our knowledge, in the pursuit of understanding the functional role of MIF in pancreatic cancer progression, the current study is the first attempt in which exogenous recombinant MIF has been used to induce high circulating MIF level in pancreatic cancer tumor-bearing animals. The results of this study suggest that MIF of non-cancer cell origin can also promote pancreatic tumor growth. Although HapT1 cells express MIF receptors, addition of exogenous MIF showing no significant effect on HapT1 cells growth *in vitro* indicates that these cells are less dependent on MIF in a growth-factor enriched culture condition (**Figure 5A&B**). A similar kind of effect of MIF has also been reported in mouse 4T1 breast tumor cell line [52]. On the other hand, overexpression of *Vegf* in HapT1 cells upon stimulation with rha-MIF and presence of more number of blood vessels in HapT1 tumors suggest a possible mechanism through which circulating MIF might indirectly enhance HapT1 tumor growth by promoting angiogenesis *in vivo*. Effect of rha-MIF in promoting the growth of HapT1 tumors corroborates and confirms the pro-tumorigenic role of MIF in pancreatic cancer progression. Hence, while designing MIF-targeted therapy in pancreatic cancer, all the possible sources of MIF that can have effect on overall tumor growth need to be taken into consideration.

Moreover, in future, a better understanding about the mechanisms through which circulating MIF can promote pancreatic tumor progression might provide a rationale to design effective therapeutic strategy against this deadly disease.

## Conclusion

In the current study, hamster MIF has been cloned, expressed and purified for the first time, which will be of immense value for future studies. The crystal structure of MIF will provide direction to study binding affinity of drugs or inhibitors for therapeutic targeting, which will facilitate experiments on the functional role of MIF in different pathological conditions in a hamster model. As we show hamster MIF to be active both enzymatically and biologically, we believe this work will lead to improved strategies for understanding this molecule in the context of different diseases, including cancer in a hamster model. Importantly, the current study provides convincing experimental evidence which suggests that hamster model of pancreatic cancer can be reliably used to investigate MIF related questions in pancreatic cancer.

## Acknowledgment

We sincerely acknowledge the support provided by the Institute of Life Sciences, Bhubaneswar to SBS and DV; extramural funding by SERB, DST, India (EMR/2016/006214) to SBS; SERB National Post-Doctoral Fellowship to PD; and DBT Research Fellowship to RS. Data collection at ESRF-ID30A-3 was facilitated by the DBT supported ESRF Access Program (BT/INF/22/SP22660/2017).

## Disclosure/Conflict of interest

The authors declare no conflict of interest.

## REFERENCES

1. Niklasson, B.S., G.F. Meadors, and C.J. Peters, Active and passive immunization against Rift Valley fever virus infection in Syrian hamsters. Acta Pathol Microbiol Immunol Scand C, 1984. 92(4): p. 197–200.

2. Milazzo, M.L., et al., Maporal viral infection in the Syrian golden hamster: a model of hantavirus pulmonary syndrome. J Infect Dis, 2002. 186(10): p. 1390–5.

3. Fisher, A.F., et al., Induction of severe disease in hamsters by two sandfly fever group viruses, Punta toro and Gabek Forest (Phlebovirus, Bunyaviridae), similar to that caused by Rift Valley fever virus. Am J Trop Med Hyg, 2003. 69(3): p. 269–76.

4. Tesh, R.B., et al., Experimental yellow fever virus infection in the Golden Hamster (Mesocricetus auratus). I. Virologic, biochemical, and immunologic studies. J Infect Dis, 2001. 183(10): p. 1431–6.

5. Xiao, S.Y., et al., West Nile virus infection in the golden hamster (Mesocricetus auratus): a model for West Nile encephalitis. Emerg Infect Dis, 2001. 7(4): p. 714–21.

6. Wong, K.T., et al., A golden hamster model for human acute Nipah virus infection. Am J Pathol, 2003. 163(5): p. 2127–37.

7. Roberts, A., et al., Severe acute respiratory syndrome coronavirus infection of golden Syrian hamsters. J Virol, 2005. 79(1): p. 503–11.

8. Bosco-Lauth, A.M., et al., Development of a Hamster Model for Chikungunya Virus Infection and Pathogenesis. PLoS One, 2015. 10(6): p. e0130150.

9. Pour, P.M., et al., Current knowledge of pancreatic carcinogenesis in the hamster and its relevance to the human disease. Cancer, 1981. 47(6 Suppl): p. 1573–89.

10. Suklabaidya, S., et al., Characterization and use of HapT1-derived homologous tumors as a preclinical model to evaluate therapeutic efficacy of drugs against pancreatic tumor desmoplasia. Oncotarget, 2016. 7(27): p. 41825–41842.

11. Ramachandhiran, D., V. Vinothkumar, and S. Babukumar, Paeonol exhibits anti-tumor effects by apoptotic and anti-inflammatory activities in 7,12-dimethylbenz(a) anthracene induced oral carcinogenesis. Biotech Histochem, 2018: p. 1–16.

12. Pal, R., et al., In-vivo topical mucosal delivery of a fluorescent deoxy-glucose delineates neoplasia from normal in a preclinical model of oral epithelial neoplasia. Sci Rep, 2018. 8(1): p. 9760.

13. Salzwedel, A.O., et al., Combination of interferon-expressing oncolytic adenovirus with chemotherapy and radiation is highly synergistic in hamster model of pancreatic cancer. Oncotarget, 2018. 9(26): p. 18041–18052.

14. Benali, N., et al., Inhibition of growth and metastatic progression of pancreatic carcinoma in hamster after somatostatin receptor subtype 2 (sst2) gene expression and administration of cytotoxic somatostatin analog AN-238. Proc Natl Acad Sci U S A, 2000. 97(16): p. 9180–5.

15. Wang, Y., et al., Experimental Models in Syrian Golden Hamster Replicate Human Acute Pancreatitis. Sci Rep, 2016. 6: p. 28014.

16. Liu, C., et al., Effects of berberine on amelioration of hyperglycemia and oxidative stress in high glucose and high fat diet-induced diabetic hamsters in vivo. Biomed Res Int, 2015. 2015: p. 313808.

17. Bifulco, C., et al., Tumor growth-promoting properties of macrophage migration inhibitory factor. Curr Pharm Des, 2008. 14(36): p. 3790–801.

18. Santos, L.L. and E.F. Morand, Macrophage migration inhibitory factor: a key cytokine in RA, SLE and atherosclerosis. Clin Chim Acta, 2009. 399(1-2): p. 1–7.

19. Kleemann, R. and R. Bucala, Macrophage migration inhibitory factor: critical role in obesity, insulin resistance, and associated comorbidities. Mediators Inflamm, 2010. 2010: p. 610479.

20. Gilliver, S.C., et al., MIF: a key player in cutaneous biology and wound healing. Exp Dermatol, 2011. 20(1): p. 1–6.

21. Babu, S.N., G. Chetal, and S. Kumar, Macrophage migration inhibitory factor: a potential marker for cancer diagnosis and therapy. Asian Pac J Cancer Prev, 2012. 13(5): p. 1737–44.

22. Micsonai, A., et al., BeStSel: a web server for accurate protein secondary structure prediction and fold recognition from the circular dichroism spectra. Nucleic Acids Res, 2018. 46(W1): p. W315–W322.

23. Kumar, S., G. Stecher, and K. Tamura, MEGA7: Molecular Evolutionary Genetics Analysis Version 7.0 for Bigger Datasets. Mol Biol Evol, 2016. 33(7): p. 1870–4.

24. Saitou, N. and M. Nei, The neighbor-joining method: a new method for reconstructing phylogenetic trees. Mol Biol Evol, 1987. 4(4): p. 406–25.

25. Till, M., et al., Improving the success rate of protein crystallization by random microseed matrix screening. J Vis Exp, 2013(78).

26. Kabsch, W., XDS. Acta Crystallogr D Biol Crystallogr, 2010. 66(Pt 2): p. 125–32.

27. Vagin, A. and A. Teplyakov, Molecular replacement with MOLREP. Acta Crystallogr D Biol Crystallogr, 2010. 66(Pt 1): p. 22–5.

28. The CCP4 suite: programs for protein crystallography. Acta Crystallogr D Biol Crystallogr, 1994. 50(Pt 5): p. 760–3.

29. Murshudov, G.N., et al., REFMAC5 for the refinement of macromolecular crystal structures. Acta Crystallogr D Biol Crystallogr, 2011. 67(Pt 4): p. 355–67.

30. Emsley, P. and K. Cowtan, Coot: model-building tools for molecular graphics. Acta Crystallogr D Biol Crystallogr, 2004. 60(Pt 12 Pt 1): p. 2126–32.

31. Laskowski, R.A., et al., PROCHECK: a program to check the stereochemical quality of protein structures. Journal of Applied Crystallography, 1993. 26(2): p. 283–291.

32. Dios, A., et al., Inhibition of MIF bioactivity by rational design of pharmacological inhibitors of MIF tautomerase activity. J Med Chem, 2002. 45(12): p. 2410–6.

33. Mookerjee, A., P.C. Sen, and A.C. Ghose, Immunosuppression in hamsters with progressive visceral leishmaniasis is associated with an impairment of protein kinase C activity in their lymphocytes that can be partially reversed by okadaic acid or anti-transforming growth factor beta antibody. Infect Immun, 2003. 71(5): p. 2439–46.

34. Sun, H.W., et al., Crystal structure at 2.6-A resolution of human macrophage migration inhibitory factor. Proc Natl Acad Sci U S A, 1996. 93(11): p. 5191–6.

35. Mitchell, R., et al., Cloning and characterization of the gene for mouse macrophage migration inhibitory factor (MIF). J Immunol, 1995. 154(8): p. 3863–70.

36. Dobson, S.E., et al., The crystal structures of macrophage migration inhibitory factor from Plasmodium falciparum and Plasmodium berghei. Protein Sci, 2009. 18(12): p. 2578–91.

37. Bozza, M., et al., Structural characterization and chromosomal location of the mouse macrophage migration inhibitory factor gene and pseudogenes. Genomics, 1995. 27(3): p. 412–9.

38. Lubetsky, J.B., et al., Pro-1 of macrophage migration inhibitory factor functions as a catalytic base in the phenylpyruvate tautomerase activity. Biochemistry, 1999. 38(22): p. 7346–54.

39. Al-Abed, Y. and S. VanPatten, MIF as a disease target: ISO-1 as a proof-of-concept therapeutic. Future Med Chem, 2011. 3(1): p. 45–63.

40. Lubetsky, J.B., et al., The tautomerase active site of macrophage migration inhibitory factor is a potential target for discovery of novel antiinflammatory agents. J Biol Chem, 2002. 277(28): p. 24976–82.

41. Sommerville, C., et al., Biochemical and immunological characterization of Toxoplasma gondii macrophage migration inhibitory factor. J Biol Chem, 2013. 288(18): p. 12733–41.

42. Crichlow, G.V., et al., Alternative chemical modifications reverse the binding orientation of a pharmacophore scaffold in the active site of macrophage migration inhibitory factor. J Biol Chem, 2007. 282(32): p. 23089–95.

43. El-Turk, F., et al., Characterization of molecular determinants of the conformational stability of macrophage migration inhibitory factor: leucine 46 hydrophobic pocket. PLoS One, 2012. 7(9): p. e45024.

44. Kudrin, A., et al., Human macrophage migration inhibitory factor: a proven immunomodulatory cytokine? J Biol Chem, 2006. 281(40): p. 29641–51.

45. Rajasekaran, D., et al., Macrophage Migration Inhibitory Factor-CXCR4 Receptor Interactions: EVIDENCE FOR PARTIAL ALLOSTERIC AGONISM IN COMPARISON WITH CXCL12 CHEMOKINE. J Biol Chem, 2016. 291(30): p. 15881–95.

46. White, D.A., et al., Proinflammatory action of MIF in acute myocardial infarction via activation of peripheral blood mononuclear cells. PLoS One, 2013. 8(10): p. e76206.

47. Calandra, T., et al., MIF as a glucocorticoid-induced modulator of cytokine production. Nature, 1995. 377(6544): p. 68–71.

48. Amin, M.A., et al., Migration inhibitory factor up-regulates vascular cell adhesion molecule-1 and intercellular adhesion molecule-1 via Src, PI3 kinase, and NFkappaB. Blood, 2006. 107(6): p. 2252–61.

49. Costa-Silva, B., et al., Pancreatic cancer exosomes initiate pre-metastatic niche formation in the liver. Nat Cell Biol, 2015. 17(6): p. 816–26.

50. Yang, S., et al., A Novel MIF Signaling Pathway Drives the Malignant Character of Pancreatic Cancer by Targeting NR3C2. Cancer Res, 2016. 76(13): p. 3838–50.

51. Richard, V., et al., Involvement of macrophage migration inhibitory factor and its receptor (CD74) in human breast cancer. Oncol Rep, 2014. 32(2): p. 523–9.

52. Simpson, K.D., D.J. Templeton, and J.V. Cross, Macrophage migration inhibitory factor promotes tumor growth and metastasis by inducing myeloid-derived suppressor cells in the tumor microenvironment. J Immunol, 2012. 189(12): p. 5533–40.

53. Chesney, J.A. and R.A. Mitchell, 25 Years On: A Retrospective on Migration Inhibitory Factor in Tumor Angiogenesis. Mol Med, 2015. 21 Suppl 1: p. S19–24.

54. Wold, W.S. and K. Toth, Chapter three--Syrian hamster as an animal model to study oncolytic adenoviruses and to evaluate the efficacy of antiviral compounds. Adv Cancer Res, 2012. 115: p. 69–92.

55. Buchko, G.W., et al., Crystal structure of a macrophage migration inhibitory factor from Giardia lamblia. J Struct Funct Genomics, 2013. 14(2): p. 47–57.

56. Rosengren, E., et al., The macrophage migration inhibitory factor MIF is a phenylpyruvate tautomerase. FEBS Lett, 1997. 417(1): p. 85–8.

57. Taylor, A.B., et al., Crystal structure of macrophage migration inhibitory factor complexed with (E)-2-fluoro-p-hydroxycinnamate at 1.8 A resolution: implications for enzymatic catalysis and inhibition. Biochemistry, 1999. 38(23): p. 7444–52.

58. Ouertatani-Sakouhi, H., et al., Identification and characterization of novel classes of macrophage migration inhibitory factor (MIF) inhibitors with distinct mechanisms of action. J Biol Chem, 2010. 285(34): p. 26581–98.

59. Calandra, T. and T. Roger, Macrophage migration inhibitory factor: a regulator of innate immunity. Nat Rev Immunol, 2003. 3(10): p. 791–800.

60. Mitchell, R.A., et al., Macrophage migration inhibitory factor (MIF) sustains macrophage proinflammatory function by inhibiting p53: regulatory role in the innate immune response. Proc Natl Acad Sci U S A, 2002. 99(1): p. 345–50.

61. Emonts, M., et al., Association between high levels of blood macrophage migration inhibitory factor, inappropriate adrenal response, and early death in patients with severe sepsis. Clin Infect Dis, 2007. 44(10): p. 1321–8.

62. Shi, C. and E.G. Pamer, Monocyte recruitment during infection and inflammation. Nat Rev Immunol, 2011. 11(11): p. 762–74.

63. Girard, E., et al., Macrophage migration inhibitory factor produced by the tumour stroma but not by tumour cells regulates angiogenesis in the B16-F10 melanoma model. Br J Cancer, 2012. 107(9): p. 1498–505.

64. Choudhary, S., et al., Macrophage migratory inhibitory factor promotes bladder cancer progression via increasing proliferation and angiogenesis. Carcinogenesis, 2013. 34(12): p. 2891–9.

65. White, E.S., et al., Macrophage migration inhibitory factor and CXC chemokine expression in non-small cell lung cancer: role in angiogenesis and prognosis. Clin Cancer Res, 2003. 9(2): p. 853–60.

66. Coleman, A.M., et al., Cooperative regulation of non-small cell lung carcinoma angiogenic potential by macrophage migration inhibitory factor and its homolog, D-dopachrome tautomerase. J Immunol, 2008. 181(4): p. 2330–7.

