## Supplementary figures and images for "Macrophage migration inhibitory factor of Syrian golden hamster has similar structure and function as human MIF and promotes pancreatic tumor growth"

### Supplementary 1

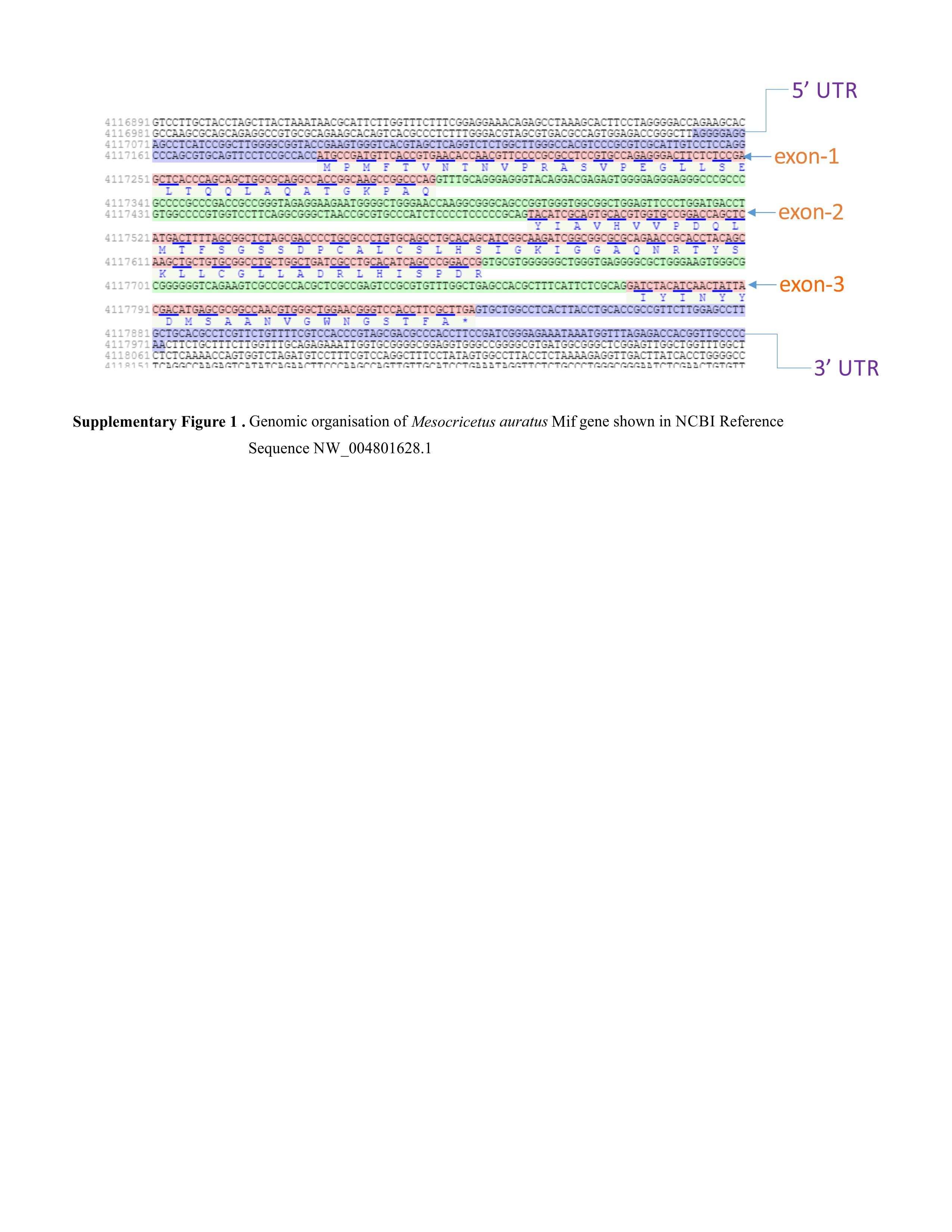

### Supplementary 2

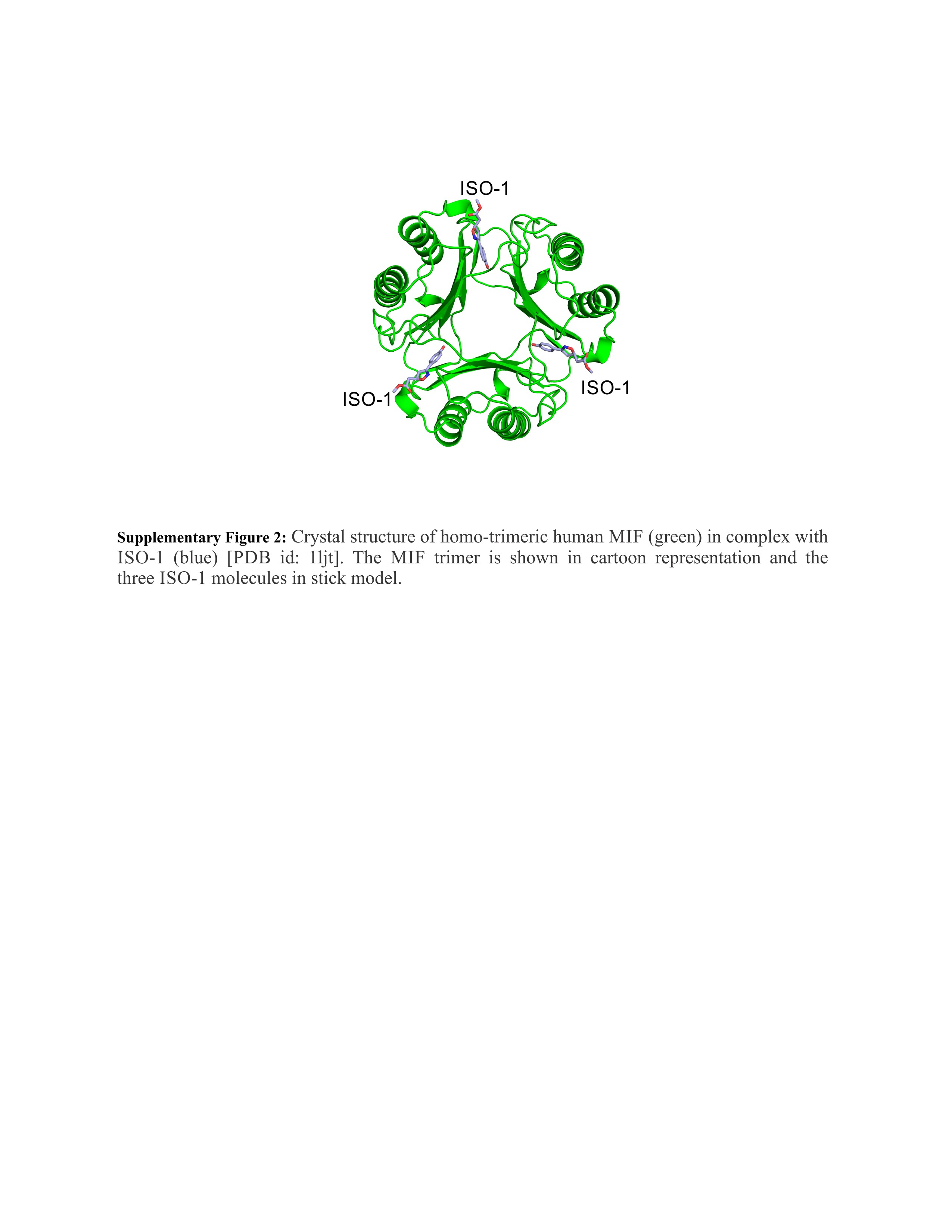

### Supplementary 3

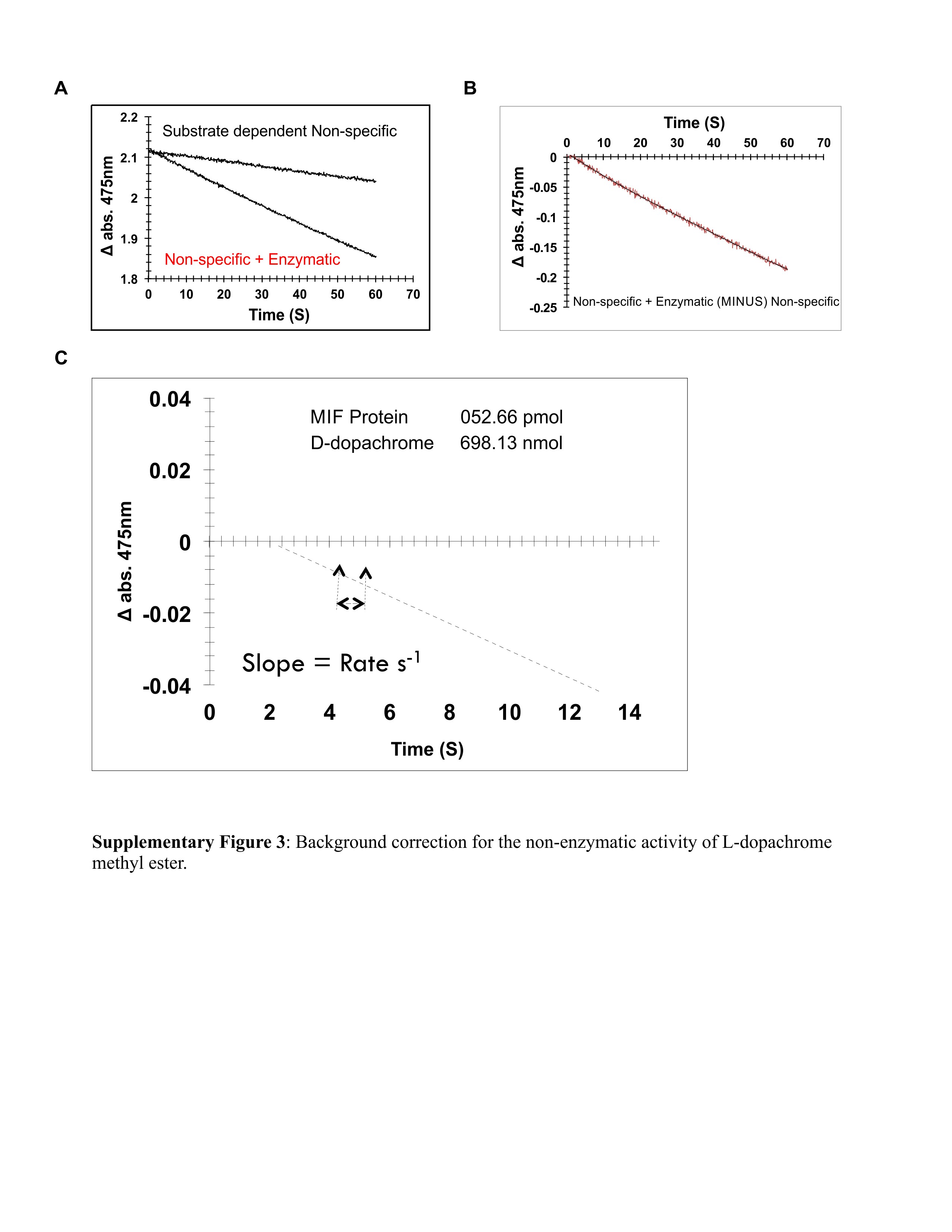
